# Assisting and Accelerating NMR Assignment with Restrained Structure Prediction

**DOI:** 10.1101/2023.04.14.536890

**Authors:** Sirui Liu, Haotian Chu, Yuhao Xie, Fangming Wu, Ningxi Ni, Chenghao Wang, Fangjing Mu, Jiachen Wei, Jun Zhang, Mengyun Chen, Junbin Li, Fan Yu, Hui Fu, Shenlin Wang, Changlin Tian, Zidong Wang, Yi Qin Gao

## Abstract

NMR experiments can detect in situ structures and dynamic interactions, but the NMR assignment process requires expertise and is time-consuming, thereby limiting its applicability. Deep learning algorithms have been employed to aid in experimental data analysis. In this work, we developed a RASP model which can enhance structure prediction with restraints. Based on the Evoformer and structure module architecture of AlphaFold, this model can predict structure based on sequence and a flexible number of input restraints. Moreover, it can evaluate the consistency between the predicted structure and the imposed restraints. Based on this model, we constructed an iterative NMR NOESY peak assignment pipeline named FAAST, to accelerate assignment process of NOESY restraints and obtaining high quality structure ensemble. The RASP model and FAAST pipeline not only allow for the leveraging of experimental restraints to improve model prediction, but can also facilitate and expedite experimental data analysis with their integrated capabilities.

## Introduction

NMR is an experimental technique used to determine structures and detect weak interactions in situ^1,2^. However, NMR assignment requires both expertise and time. It might take months even years for NMR assignment. Leveraging machine learning and deep learning technologies, researchers have endeavored to automate the NMR assignment protocol. For example, ARTINA^3^ provides an integrated pipeline which accepts raw NMR spectra, assigns chemical shifts and NOE peaks, and provides structures simultaneously. It utilizes molecular simulations to construct structures, with a focus on achieving automated and accurate chemical shift assignments. Specifically, the NOESY peak assignment process provides hydrogen restraints and is an essential technique in NMR structure analysis, although the structure construction mostly relies on molecular simulation. Automated algorithms such as CYANA^4^, ARIA^5,6^, CANDID^7^ have been developed to assist NOE peak assignment, which mostly apply strategies such as molecular dynamics simulation or simulated annealing for structure construction, thus is relatively time consuming. Rosetta suites or pipelines leveraging sparse NMR restraints from NOE, RDC, and PRE data have also be developed for the data-assisted structure construction^8,9,10^.

Recent progress in deep learning provides more efficient and accurate tools to generate protein structure given its sequence. Employing deep learning protein structure prediction models for experimental data analysis has been a problem of interest. In practice, the input data form and distribution of general structure prediction models, such as AlphaFold^11^, do not necessarily align with the needs of experimental methods. While AlphaFold and AlphaFold-multimer^12^ have greatly improved the accuracy of predicting static protein structures, unresolved issues remain, such as generating dynamic structures and predicting restrained structures. Questions remain on how experimental information can facilitate rapid structure prediction and how structure prediction methods can aid in the resolution or acceleration of experimental data analysis. Attempts have been made to provide AlphaFold structures as templates for X-ray replacement^13^ or Cryo-EM density map^14^ templates. However, these approaches rely on iterative template use, which includes dense but not necessarily accurate restraints and cannot utilize structural differences to improve predictions. Recently, AlphaLink^15^ fine-tuned AlphaFold to accept sparse restraints, improving AlphaFold performance in cross-linking experiments.

In this work we propose a model named Restraints Assisted Structure Predictor (RASP) and an iterative NMR NOESY peak assignment pipeline called FAAST(iterative Folding Assisted peak ASsignmenT). The architecture of RASP is derived from AlphaFold Evoformer and structure module, and it accepts abstract or experimental restraints, sparse or dense, to generate structures. This enables RASP to be used in diverse applications, including improving structure predictions for multi-domain proteins and those with few multiple sequence alignments (MSAs). The confidence of RASP can evaluate restraint quality in terms of information efficiency and accuracy. Consequently, by leveraging the model’s ability to accept a flexible number of restraints and evaluate them, together with an NMR assignment protocol adapted from ARIA^6^, we developed the FAAST pipeline. Using chemical shift and NOE peak lists as input, FAAST assigns NOE peaks iteratively and generates a structure ensemble based on the subsampled restraints, thus accelerating NMR analysis.

## Results

### RASP takes in restraints directly and helps in structure prediction

To facilitate general experimental information as restraints, we developed the RASP model based on the AlphaFold architecture. The model takes in sequence and a flexible number of distance restraints and returns a structure that largely complies with the restraints. Additionally, it measures the consistency between the restraints and sequences. We consider restraints as a form of edge information and use the edge bias in the Evoformer MSA attention block (defined as MSA bias) and invariant point attention block (IPA, defined as IPA bias) (Fig. 1a, 1b). Moreover, we experimented with the pair representation update in Evoformer (defined as pair bias) and adopted the structure module (defined as structure bias) to ensure that the structure follows the restraints in one update. We evaluated the impact of the four types of bias information and chose MSA and IPA biases as the baseline RASP model setting for simplicity and stability (Suplementary Figure 1). We use only the first two bias forms in the following experiments.

**Fig. 1.**
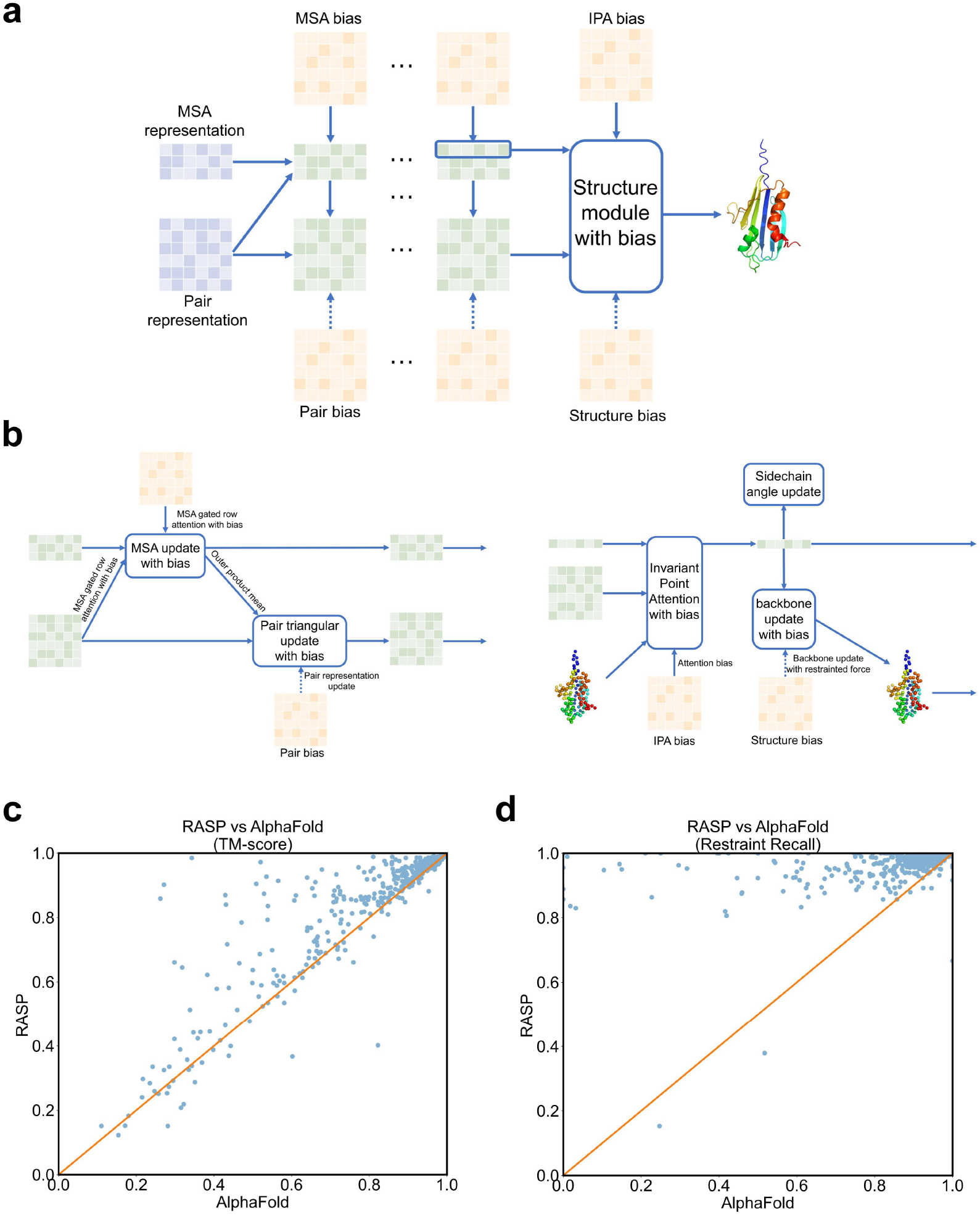
RASP model takes in restraints in different forms and outperforms AlphaFold with restraint assistance. **a**. A scheme of how the pairwise restraints can be taken into the model in the form of MSA, pair, IPA, and structure biases, adopted from the Evoformer and structure module in AlphaFold. **b**. More detailed illustration of the model structure with restraints treated as biases in the Evoformer block (left) and structure block (right). For the PSP validation dataset, RASP with a fixed number of randomly sampled input restraints outperforms AlphaFold on **c**. the TM-score between the PDB structure and predicted structure and **d**. on restraint recall.

We implemented RASP using MindSpore^16^ and trained it on 32 * Ascend910 NPUs. We initialized the model with MEGA-Fold^17^ weights and fine-tuned it on the PSP dataset^17^ with a true structure: distilled structure ratio of 1:3. We sampled pairwise restraints to tolerate distance noise (refer to Methods). The training converged after 15k steps (480k samples in total, Supplementary Figure 2) and demonstrated stable improvement over initial MEGA-Fold.

Although the model supports templates in prediction and could improve performance with template used, for fairness and to avoid data leakage, we chose not to utilize templates in this research. We tested the model’s performance on the PSP validation dataset^17^ previously constructed along with the PSP training set, which contains 490 samples of CAMEO^18,19,20,21^ targets and unique proteins between October 2021 and March 2022. This validation set is strictly after the PDB and sequence deposition time of training set. When restricting the number of restraints to 100, the TM-score^22,23^ which measures topological similarity between structures improved significantly for the structure prediction in the PSP validation dataset (Fig. 1c). Furthermore, the model followed the randomly sampled restraints much better than those predicted by AlphaFold or MEGA-fold, as expected (Fig. 1D, Supplementary Figure 3). Moreover, the violation loss which measures bond length, bond angle, and atomic clash violation for RASP predictions remained low with a median of 0.0012 (Supplementary Figure 4), indicating its capability to predict structures following basic physiochemical principles.

### RASP helps structure prediction and evaluation in a broad range of restraints numbers

We discovered that the structure accuracy improves steadily as the number of restraints increases, starting from zero (Fig. 2a). However, restraints recall remains relatively constant, implying that the current model can tolerate different numbers of prior information or restraints without adversely affecting the baseline model performance (Fig. 2b). Additionally, the predicted local-distance difference test score (pLDDT score) of the model serves as an indicator of the model’s confidence in the restraints. For proteins with varying numbers of restraints, the pLDDT score rises stably though not significantly with an increase in the number of restraints applied. The pLDDT confidence correlates well with the corresponding structure TM-score with an overall correlation of 0.68 (Fig. 2C).

**Fig. 2.**
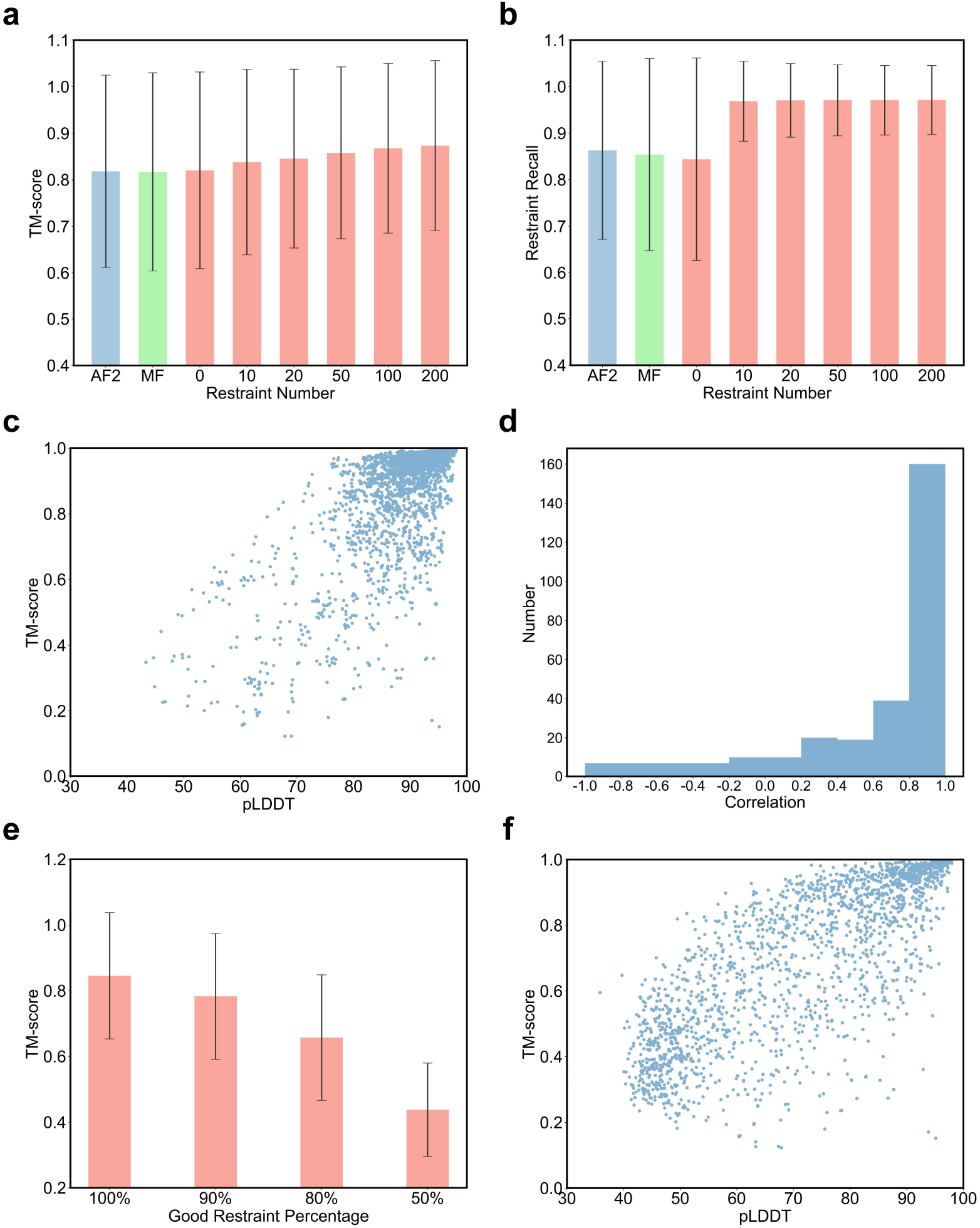
RASP is robust against restraint number and the confidence score can be used to evaluate restraint quality. Errorbars represent standard deviations for each group of data for all applicable figures. **a**.The TM-score of predicted structure by RASP steadily increases over both AlphaFold and MEGA-Fold, with increasing number of input restraints. **b**. Meanwhile regardless of the restraint number, the restraint recall is steady above that with no restraints applied for AlphaFold, MEGA-Fold, and RASP with zero restraints. The restraint recall for those without input restraints is calculated with a fixed number of randomly sampled contacts by the same sampling strategy as input restraints. **c**. The pLDDT with different restraints number and information quality correlates well with the real TM-score of the predicted structure and deposited PDB structure, with an overall correlation coefficient of 0.68. **d**. The distribution of correlation coefficient between the predicted pLDDT confidence and TM-score with different number of restraints in use shows consistency between predicted and real structure quality. The distribution is drawn for validation samples with at least one bad prediction with TM-score<0.8, since we are more interested in the model ability to distinguish bad or inefficient restraints information from the good ones than to tell the best from a bunch of very good structures with efficient restraints. The pLDDT confidence and TM-score are largely positively correlated with a median of 0.62, suggesting that pLDDT score can be regarded as an indicator how much additional information input restraints provide. **e**. The TM-score decreases with decreasing percentage of good restraints, when total input restraints number is fixed at 20. **f**. The pLDDT confidence still correlates well with the TM-scores when bad restraints are present, with an overall correlation coefficient of 0.72, demonstrating the RASP model ability in telling bad restraint information and reporting this at the same time.

Despite the difference in restraint numbers, due to sampling randomness, some restraints might provide repeating information with MSAs (and potentially templates, although templates is not used here for fairness), and in reality the restraints information provided by experiment may not be free of error. The restraint quality therefore could be considered in two aspects: one is how much additional information it provides aside from that provided by MSA and templates, another is how accurate the restraint information is. To examine the ability of the confidence score to distinguish good and bad predictions for the same protein with different additional restraints information, we first examined the confidence-TM-score correlation for proteins with at least one prediction of lower quality (defined as TM-score lower than 0.80). The average correlation score is 0.62 (Fig. 2d). Since we take only MSA and restraints as input in the benchmark, while MSA is kept the same for predictions of the same protein, better structures can therefore be attributed to more effective restraint information. This indicates that the pLDDT score can largely be used to distinguish better structures and better corresponding restraints. Furthermore, when incorrect restraints are intentionally used (restraints with a Cβ distance greater than 12 Angstrom, as defined), the TM-score decreases significantly along with the increasing incorrect ratio and fixed number of 20 restraints (Fig. 2e), suggesting that the model is sensitive to inconsistent restraints and can distinguish corresponding bad structures. The TM-scores correlate well with the pLDDT scores with an overall correlation of 0.72 (Fig. 2f, Supplementary Figure 5). These findings indicate that the pLDDT score can gauge how well the restraints may assist in structure prediction and the restraint’s quality or self-consistency, both with restraints that may be of little use and with bad restraints present. With this evaluation, the model may find applications in areas such as NMR determination (see section below).

### RASP improve structure prediction assisted by pseudo and NMR restraints

By incorporating restraints, the model demonstrates improved capability to predict the structures of multidomain and few-MSA proteins. Two cases representative of this improvement in the PSP validation dataset are 6XMV and 7NBV(Fig. 3a, 3b). 6XMV is a multi-domain protein that exhibits wrongly predicted relative domain positions by both AlphaFold and Mega-Fold. However, utilizing randomly sampled restraints corrects the inter-domain positions. For 7NBV, which is a virus protein and only has three sequences in its multiple sequence alignment, an increase in the number of randomly sampled restraints leads to a stable improvement in structure quality, with 50 restraints being used. These outcomes demonstrate the potential for using restraints to aid in the prediction of few-MSA and multi-domain proteins.

**Fig. 3.**
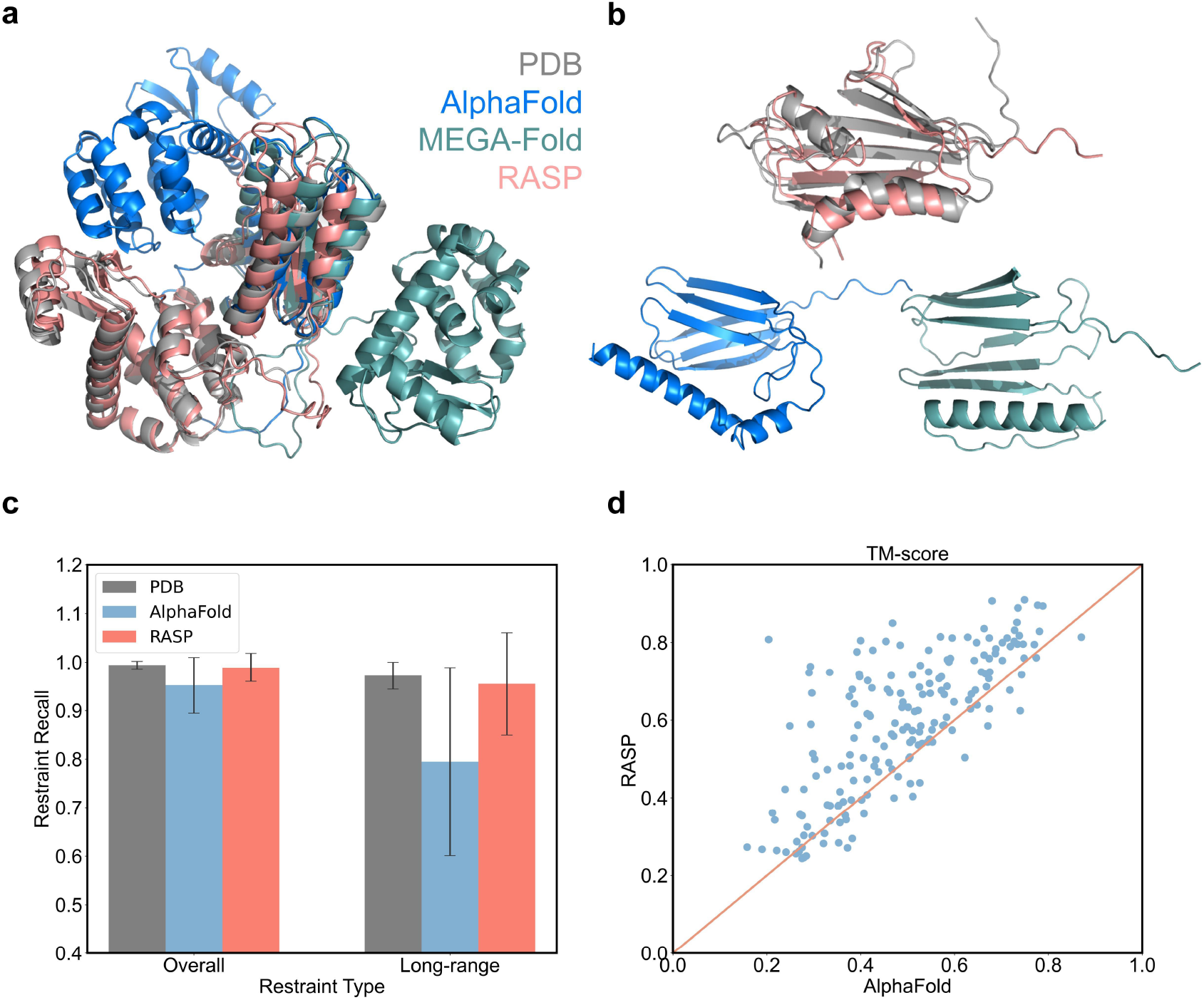
Input restraints can assist RASP prediction for multi-domain proteins, few-MSA proteins, and NMR structure prediction. **a**. For multi-domain structure 6XMV, both AlphaFold and MEGA-fold provide inaccurate relative domain positions, however with restraints RASP is able to fix the inter-domain structure. **b**. For 7NBV with few MSAs, restraint assisted prediction by RASP helps to improve the structure prediction with more accurate secondary structure and relative position between the helices and beta-sheets The structures are presented separately because the RMSDs for AlphaFold and MEGA-Fold predictions are higher than 10 Angstrom (19.0 and 15.9, respectively) and are hard to align to the PDB structure. **c**. With varied number of deposited .mr restraints provided by the PDB databank, NMR structures that fail for AlphaFold prediction can be fixed in terms of overall and long-range (sequence separation≥4) restraint recall and **d**. TM-score that measures the similarity between the predicted structure and the deposited ones.

NMR is a commonly used experimental structure determination method that generates restraints of different magnitudes. Despite that in many cases AlphaFold predictions follow the restraints similar or even better than deposited NMR PDB structures^24,25^, AlphaFold does not naturally foresee structures compatible with NMR restraints and may produce alternative structures as opposed to those deposited in the PDB. Given the continued evolution of NMR data deposit requirements and the length of time during which samples may be deposited, some entries in the PDB and BMRB database do not include restraint files. After filtration of NMR samples deposited in the RCSB PDB bank with restraint files (.mr) available and bad AlphaFold predictions (long-range restraint recall lower than 90%), 182 samples remain (Supplementary Table 1). The samples exhibit a wide variation in the overall number of NMR restraints, from tens to thousands, with a median restraint quantity of 11.4 per residue. When leveraging NMR restraints to aid in structure prediction, the predicted structures better adhere to the restraints than AlphaFold predictions, both for overall restraints and especially for long-range restraints (defined as sequence separation≥4 in this work), with median restraint recall increasing from 95.2% to 99.2% and 79.5% to 96.2%, respectively (Fig. 3c). The structures generated by RASP are interestingly more consistent with the deposited structures (Fig. 3d).

### NMR NOESY assignment pipeline FAAST

With the ability of the RASP model to take restraints from a wide range of sources and evaluate their quality with pLDDT scores, it has the potential to accelerate NMR NOESY peak assignments. These assignments accumulate over assigning iterations - starting with only a few correctly identified restraints - and lead to refined structure predictions. By combining the RASP model with the ARIA^6^ assignment protocol, we built an iterative NMR analysis pipeline named FAAST (Fig. 4a). FAAST takes chemical shift and NOE peak lists as input and outputs peak assignment and structure ensembles. Each iteration involves subsampling the assigned restraints with an increasing ratio from the previous iteration as RASP input and generating an ensemble of 20 structures, which is then used for the subsequent NOE peak assignment. As pLDDT scores reflect the restraint quality, if the median pLDDT of the ensemble is lower than 80, we restart the second round of iteration with a lower restraint subsampling ratio to reduce restraints conflict (Fig. 4b). The protocol allows for a maximum of one restart, resulting in a total ensemble iteration number of 2 or 5.

**Fig. 4.**
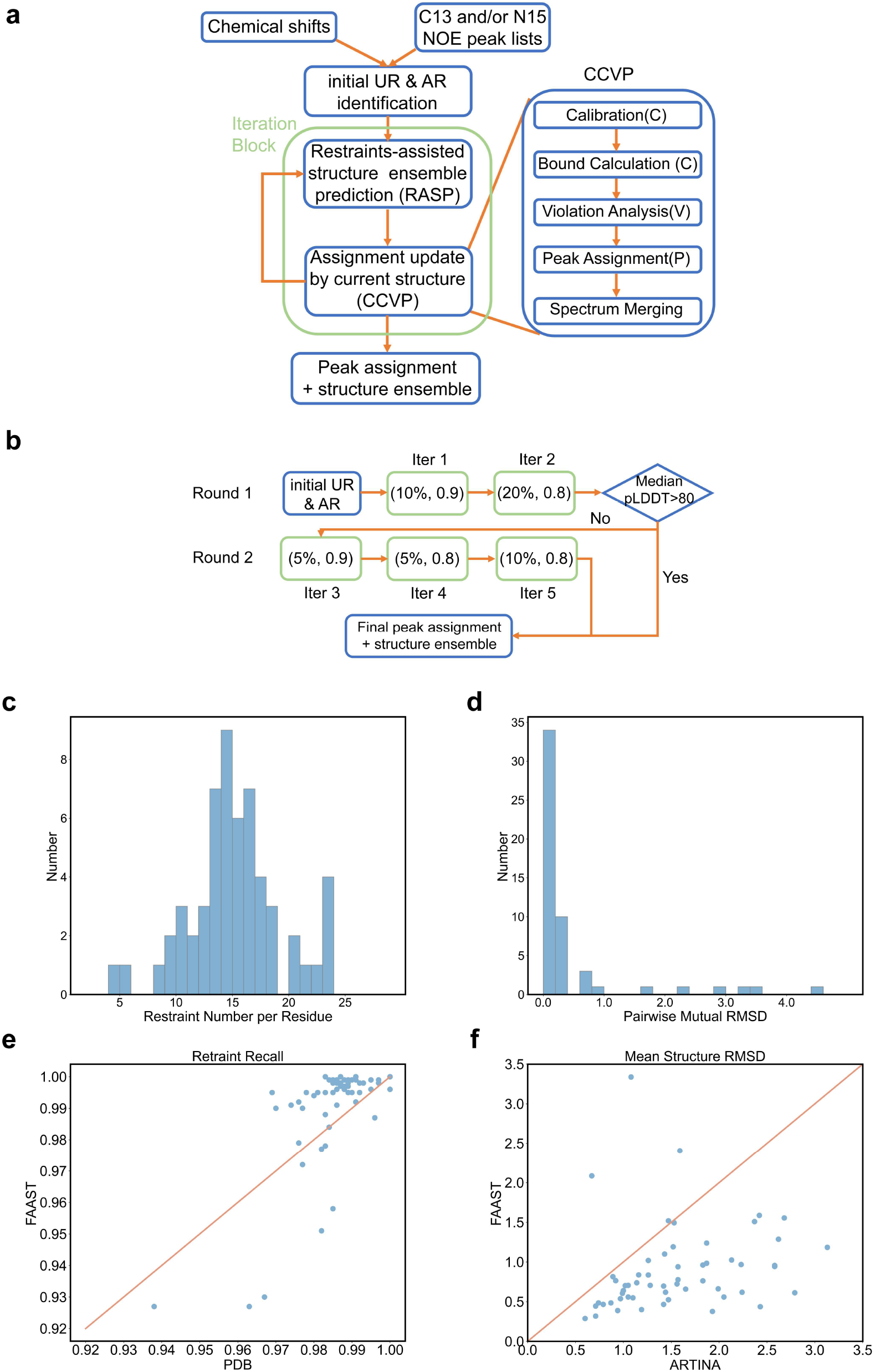
The FAAST NOESY assignment pipeline provides fast and accurate structure ensemble and NOE peak assignment at the same time. **a**. The schematic figure of the FAAST workflow. **b**. The current assignment pipeline and parameters applied. In each iteration block, the first parameter is the percentage of input restraints subsampled for RASP structure prediction in order to construct a structure bundle, the second parameter is the cumulative probability to rule out ambiguous restraints in assignment, which is also adjustable in ARIA. **c**. The distribution for number of NOE assignments per residue for the 57 ARTINA samples. The median number is around 15 with peak lists from the BMRB database. **d**. The distribution for pairwise mutual RMSD for C-alpha atoms of the structure bundle. The median pairwise RMSD is 0.87, indicating the predicted structures within each bundle are close to each other. **e**. The restraint recall is compared between the FAAST structure with the identified restraints and the deposited PDB with the deposited NMR restraints. The solved FAAST structures better follow the NMR restraints than the PDBs. **f**. The FAAST structures also exhibit lower backbone RMSD than those reported by ARTINA.

We benchmarked the FAAST pipeline on samples used in ARTINA. Out of the 100 ARTINA samples, only 57 had both chemical shift and at least one 3D NOESY peak list deposited on BMRB and can be identified from the nmrstar files (Supplementary Table 2). We validated the NMR pipeline on all of the 57 samples. With a median time of 32 minutes (minimum and maximum time of 14 and 103 minutes), we were able to assign a median of 1569 peaks per sample and a median peak number of 14.75 per residue (Fig. 4c). Furthermore, the average pairwise mutual C-alpha RMSD for the structure ensemble is 0.87 (Fig. 4d), indicating consistency between subsampled restraints and the resulted structure ensemble. We note here that pairwise RMSDs from the structure bundle in the initial iteration have a median 1.99, higher than that from the final structures, indicating that the subsampling strategy is able to generate diversified structures, and that iterative refinement leads to convergence in structure ensemble.

Moreover, the predicted structure is consistent with simultaneously assigned restraints as well as the NOE peaks. A median of 99.6% of the identified restraints match the highest confidence structure and the corresponding median is 99.0% for identified long-range restraints. In comparison, the model 1 structure and restraints from the PDB database conform on a median of 98.6% and 98.2% for all restraints and long-range restraints, respectively (Fig. 4e, Supplementary Figure 6). The RMSD score and correlation score calculated by ANSURR^26^ and DP score by RPF^27^ indicate that the structures obtained by FAAST are of comparable or better quality and consistency between predicted structure ensemble and the NOE peak lists, compared to corresponding PDB structures (Supplementary Figure 7).

The predicted structures not only agree with the assigned peaks but are also consistent with the deposited restraint and structure data. 96.9% of the deposited restraints from PDB database align with the predicted structure ensemble. The median mean structure backbone RMSD against the deposited PDB model 1 is 0.739 Angstrom for structured regions defined by ARTINA. For the median scored structure in the structure ensemble, the backbone RMSD is 0.791 Angstrom against the PDB structure. Both are lower than that reported by ARTINA, in which the median mean structure backbone RMSD is 1.44A for all samples and 1.47A for the 57 samples with BMRB peak lists (Fig. 4f).

Since only processed NOESY peak lists are available and raw peak lists absent for samples downloaded directly from BMRB, we validated the pipeline’s performance on the 2MRM case with raw NOESY peak lists. For this YgaP protein, much more restraints can be assigned from the raw peak lists than from the deposited NOE peak lists (14.19 per residue for raw lists compared to 4.93 for deposited ones). The number of assigned long-range restraints is also higher (366 for raw lists and 285 for deposited lists). Despite similar small mutual RMSDs, the predicted structures from raw peak lists and deposited peak lists have similar TM-scores to the deposited structure (0.862 and 0.857, respectively), even though the restraints assigned from the raw peak list are in better consistency than those assigned from deposited lists, with the former has a restraint recall of 98.2% and the latter of 94.9%. These results indicate that the pipeline doesn’t require strict peak assignment, and we expect the raw peak lists from NOE spectra to provide better assignment in the FAAST pipeline than the deposited peaks.

In summary, we have presented a fast NMR pipeline that provides accurate structure ensemble and highly consistent NOE peak assignments. Compared to previous methods, this pipeline is fast, and through restraints iteration and subsampling, can provide a structure ensemble plus a full set of NOE peak assignments. We expect this FAAST pipeline to be useful in the NOESY peak assignment and NMR structure determination since it performs well both with raw peak lists and deposited ones, even better with raw peak lists for the example case in this study.

## Discussion

The question of how experimental results and AI methods can mutually benefit each other has been a topic of discussion, particularly with the emergence of advanced biochemical deep learning models. Here, we present the RASP model and the FAAST pipeline, wherein the former utilizes prior knowledge or restraints to improve in silico structure predictions, while the latter employs the former’s flexible number restraint-taking capability and evaluation of restraint-structure quality to accelerate NOESY peak assignment. This model and pipeline underscore the self-consistency of the two questions as an AI method capable of being assisted by external knowledge has the potential to facilitate the acquisition and/or validation of that external knowledge in return.

Despite the application of the RASP model on NMR restraints, due to its improvement of structure prediction with abstract randomly sampled restraints, flexibility in restraint number and ability to evaluate restraints, we expect it to be useful for broad knowledge types, such as cross-linking or covalent labeling data as in AlphaLink, even abstract prior knowledge such as closeness of two residues regardless of the knowledge source, and in this way may help the generation of dynamic structures or states guided by restraints.

While we have currently applied our standard pipeline in FAAST for benchmark, parameters for RASP and CCVP steps can be flexibly adjusted by users to accommodate their particular peak quality and expectations on peak-structure convergence. When benchmarking the pipeline, we did not employ parallel computation considering the possibly limited computational resources for users. However, both the RASP prediction and relaxation can be executed parallelly, which is expected to accelerate the process up to 20 times, which is the ensemble size, depending on the hardware available. Since the chemical shift and NOE peak assignment could be iteratively improved, merging the chemical shift and peak assignment pipelines is also expected to produce more comprehensive and accurate NMR protein assignment pipelines.

Moreover, in this study we only used restraints generated from 3D NOESY spectra, but the current pipeline could be readily expanded to other NMR data types such as 4D NOESY spectra, as long as the experimental data could be formatted as pairwise restraints. More diversified forms of experimental data also exist that might provide information for different molecule types, such as NMR for protein-small molecule interactions. In addition to the conventional paired restraints, we also expect to incorporate additional information forms(e.g. torsion angle and PRE in NMR) into our structure prediction and to develop multimer and interface prediction models. These restrained structure prediction models hold the potential to introduce an alternative approach to restrained design.

## Methods

### Structure of RASP

To incorporate restraint information, we developed the RASP model derived from the AlphaFold Evoformer and structure module. Four additional biases were added, which draw on restraint information: pair bias, MSA bias, IPA bias, and structure bias. To handle inter-residue restraints as edge information, the first 3 biases are introduced as edge biases to the Evoformer and IPA modules. This inter-residue information can be naturally converted into features of shape (*N*_*res*_, *N*_*res*_, *C*_*channels*_), similar to the pair activation in the Evoformer module of the original model. Following the strategy of merging pair activation and MSA activation in the Evoformer module, an extra contact bias is added to the row-wise attention and the outer-product mean module by pair bias. The merging of inter-residue information and per-residue information also occurs in the Invariant Point Attention. The contact information is added to the IPA attention weight matrix as IPA bias in the same way as the MSA bias. In addition to the biases in the attention, an additional bias is introduced in the structure generation process. When generating the 3D structure, near-residue pairs identified by restraint information are moved into close distances, whereas the rest of the residue pairs connected to the pairs are then moved accordingly by optimizing the inter-residue distance in the violation loss of AlphaFold. For simplicity and stability, only the RASP model with MSA bias and IPA bias are used for result analysis.

### Restraint loss and tasks

All AlphaFold losses are retained, including the auxiliary loss from the Structure Module (a combination of averaged FAPE and torsion losses on the intermediate structures), averaged cross-entropy losses for distogram and masked MSA predictions, model confidence loss, experimentally resolved loss, and violation loss.

We introduce restraint loss into the model training to reinforce the input restraint information in the final prediction. This loss comprises three components, each corresponding to a restraint-related task. The first task is a 0/1 classification task with a loss called contact classification loss. In this task, residue-wise distogram prediction of input restraints is computed, and reorganized into 2 classes (whether or not the contact exists) with cross entropy calculated using the ground truth label. The second task is to minimize the distance RMSD difference of input restraints using a loss called dRMSD contact loss. The last task is to make local structures similar to the ground truth structures, and takes a reduced version of backbone FAPE loss called contact FAPE loss, in which the errors of all atom positions are calculated in the local backbone frames of all residues in the restraints. The contact FAPE loss and dRMSD contact loss are weighted equally at 0.5 so that the three losses are of the same order of magnitude at the beginning of training. We clip the sum of the last two losses by 1.5 to avoid training clashes in abnormal training examples.

### Sampling strategy

The model was trained using the PSP dataset^17^, which was previously constructed by us. The PSP dataset is a compilation of true and distilled protein structures, and it includes sequence, structure, template, and MSA data for each protein sequence. Training data for RASP are sampled with replacement from both the true structure and distillation datasets and mixed in a ratio of 1:3.

To simulate the restraints observed in real experiments, the residue-wise distance map of the protein structure is computed using the pseudo-Cβ atom position of the residue, where the pseudo-Cβ atom is the C α atom position for glycine and Cβ for other amino acids. The restraints are sampled based on a probability distribution that decreases with residue-wise distance. When the distance is <7 Angstrom(A), the probability is equal, and it decays exponentially from 7A to 10A. With this distribution, 90% of the sampled restraints are at a distance less than or equal to 8A, and 10% of the restraints are at a distance greater than or equal to 8A. This setting provides the model with a tolerance to restraints of poor quality. The number of restraints is also randomly sampled from a distribution with equal probability for 16-128A and an exponential decay from 128A to 2048A. The expected value of this distribution is 115 Angstrom.

### NMR NOE assignment pipeline

The assignment pipeline used in this study was based on ARIA 2.3^6^ (Ambiguous Restraints for Iterative Assignment), which was developed with Python 2 by Institut Pasteur. The FAAST assignment pipeline refered to part of the ARIA method, mainly the Calibration - CalculateBounds - ViolationAnalysis – Partially (CCVP for simplicity) functions, these functions perform assignment of peaks by comparing distance of restraints atom pairs in reference structure and theoretical distance calculated from intensity volume of the peaks. The original Python 2 code is first simplified and translated to Python 3 to cooperate with other parts of FAAST. Also, as the protein structure predicted by RASP does not distinguish equivariant hydrogens in amino acids, we collected equivariant groups of 20 common amino acids and redesigned the CCVP assignment algorithm based on distances between equivalent atomic groups according to the equivariant groups list.

The initial assignment is performed by comparing the chemical shift and NOE lists. Most of the restraints generated by initial assignments are ambiguous restraints (ARs), that is, a single peak is assigned with more than one possibility. While some peaks are naturally unambiguous (URs). The quality of the initial URs could be very low, with more than half exceeding a distance of 6.0 Angstrom. Thus, for the initial assignment, we filtered out initial URs with distances larger than 12 Angstrom in the reference prediction without restraints. For each iteration, the URs are fed into RASP to generate 20 structures with the UR subsampling rate of 5%, 10%, or 20%, depending on the iteration step, and the structures are relaxed by OpenMM^28^. The structure bundle is then used to assign NOE peaks by CCPV.

In the standard pipeline, the hyper-parameters used for restraint subsampling and CCVP are iteratively tightened, with a subsampling rate of 10% and partial assignment cumulative acceptance of 0.9 for the first iteration and 20% and 0.8 for the second iteration. ARs are transformed into URs iteratively. If after the second iteration, the median pLDDT score is lower than 80, a second round of iteration is initiated with subsampling and CCVP parameters of (5%, 0.9), (5%, 0.8), and (10%, 0.8). The entire process takes 2-5 structure generation iterations, and the number of iterations, as well as the iterating parameters, can be flexibly adjusted.

### Benchmarking data

The benchmarking data for our method consist of three parts:

#### PSP validation dataset

This dataset is the validation set of the PSP dataset and is used to evaluate the performance of the RASP model. The restraints in this dataset are sampled in the same way as during model training.

#### MR dataset

For most NMR structure in RCSB PDB database^29^, restraint .mr files are also deposited. We selected all the NMR .mr files from the RCSB PDB database in which a) the restraint numbering followed the PDB numbering, and b) the restraint recall for long-range restraints of the structure predicted by AlphaFold is less than 90%. This resulted in 182 samples.

#### NMR dataset

The NMR dataset was obtained to evaluate the NMR FAAST protocol. We obtained the .star file (including the chemical shift and NOE list), .mr file (submitted restraints), and .pdb file (structure) for 100 sequences in the ARTINA dataset by crawling the BMRB^30^ and RCSB PDB databases. After filtering out the .star files with missing chemical shift or NOE lists, 57 sequences were available for testing our protocol.

Additionally, as the NOE list in .star files from the BMRB dataset before submission could be filtered, we used the raw NOE peak list for pdb id 2MRM to evaluate the peak quality.

### Evaluation methods

We evaluated the structures and their consistency with restraints mainly with TM-score, root mean square deviation (RMSD), and restraint recall.

TM-score^22,23^ is a metric for assessing the topological similarity of protein structures. This score falls between 0 and 1, and higher TM-score indicates higher similarity between the two compared proteins. We used the TM-align^31^ package downloaded from Zhang lab for calculation of TM-scores.

In FAAST evaluation, two types of RMSD calculations are used. For measurement of mutual similarity within a structure ensemble, we calculate the pairwise Cα RMSD between all pairs of different structures within the bundle, and average them to obtain the pairwise mutual RMSD. For measurement of structure similarity between the deposited PDBs and processed structure ensemble, we follow the ARTINA^3^ evaluation and calculated the mean structure backbone atom RMSD for structured regions defined by ARTINA. All RMSD calculations are performed using PyMOL.

Restraint recall is used to measure the consistency between a structure and a set of restraints. It is defined as the ratio between the number of rightly followed restraints by the structure and the number of ground truth restraints from PDB database, similar to the definition of recall in the machine learning field. In RASP evaluation, since the restraints are at residue level, we define a pairwise restraint to be followed by the structure as the distance between pseudo-Cβ atoms (see sampling strategy in method) in the residue pair is closer than 8 Angstrom. In FAAST pipeline evaluation, since the NMR restraints are at atomic level, we define a pairwise restraint to be followed only when the closest hydrogen atomic distance in the two equivalent groups from the structure is lower than 6 Angstrom.

We further evaluated the goodness-of-fit of our predicted structures by FAAST to the experimental data using correlation score, RMSD score, and DP score. ANSURR^26^ (v2.0.55) (https://github.com/nickjf/ANSURR2) was used to calculate the correlation score and RMSD score. ANSURR accesses the accuracy of query structures by comparing their local rigidity with the random coil index (RCI). Both correlation score and RMSD score fall between 0 and 100, with higher scores indicating higher accuracy of structures in the aspects of secondary structure and overall rigidity, respectively. We re-referenced chemical shifts before calculating RCI by specifying “-r” as recommended and ran with “ansurr -p xxxx.pdb -s xxxx.str -r” for each structure.

The discrimination power (DP) score is the final output of the NMR structure quality assessment web-server tool RPF^27^ (https://montelionelab.chem.rpi.edu/rpf/), implying the correctness of the overall fold of query structure. We ran RPFs in batch using “dpsimple” from ASDP (v2.3) (https://github.rpi.edu/RPIBioinformatics/ASDP_public).

## Data availability

The training set and PSP validation dataset are from our previous work and have been publicly available at http://ftp.cbi.pku.edu.cn/psp/. The PDB ID of the 182 samples used for restraints analysis in this work are available in Supplementary Table 1 and the PDB and restraint .mr files can be downloaded at RCSB PDB database(https://www.rcsb.org/). The information of the 57 samples used for FAAST pipeline benchmark are provided in Supplementary Table 2, and the structure .pdb files, restraint .mr files, and NMRSTAR .str files are available at RCSB PDB(https://www.rcsb.org/) and BMRB(https://bmrb.io/) databases, according to their PDB and BMRB entry IDs.

## Code Availability

The RASP and FAAST code are available at [our gitee repository](https://gitee.com/mindspore/mindscience/blob/master/MindSPONGE/applications/research/FAAST/README.md) under Apache 2.0 license. We additionally provide a [colab notebook](https://colab.research.google.com/drive/1uaki0Ui1Y_gqVW7KSo838aOhXHSM3PTe?usp=sharing) for ease of use.

## Author contributions

S.L., Z.W., and Y.Q.G. developed overall concepts in the paper and supervised the project. S.L., H.C., and Y.X. wrote the initial draft of manuscript. S.L., H.C., N.N., C.W., J.W., J.Z., M.C., J.L., and F.Y. developed and validated model and pipeline. S.L., H.C., Y.X, F.W., F.M., J.W., H.F., S.W., and C.T. carried out the data processing and analyses. Specifically, F.W. and C.T. provided the raw YgaP NOESY peak data. All authors contributed ideas to the work and assisted in editing of the manuscript.

## Acknowledgements

The authors thank Yupeng Huang for helpful discussions on data processing, and would like to extend our gratitude to Yuanpeng Janet Huang, the author of RPF, for his patience and guidance on how to use dpsimple. This work was supported by National Key R&D Program of China (2022ZD0115001), National Natural Science Foundation of China (92053202, 22050003, 22274050, and 21825703), the Strategic Priority Research Program of Chinese Academy of Sciences (XDB37000000), and Collaborative Innovation Program of Hefei Science Center, CAS (2022HSC-CIP011). A portion of this work was performed on the Steady High Magnetic Field Facilities, High Magnetic Field Laboratory, CAS.

